# Structure and dynamics of the SARS-CoV-2 envelope protein monomer

**DOI:** 10.1101/2021.03.10.434722

**Authors:** Alexander Kuzmin, Philipp Orekhov, Roman Astashkin, Valentin Gordeliy, Ivan Gushchin

## Abstract

Coronaviruses, especially SARS-CoV-2, present an ongoing threat for human wellbeing. Consequently, elucidation of molecular determinants of their function and interaction with host is an important task. Whereas some of the coronaviral proteins are extensively characterized, others remain understudied. Here, we use molecular dynamics simulations to analyze the structure and dynamics of the SARS-CoV-2 envelope (E) protein (a viroporin) in the monomeric form. The protein consists of the hydrophobic α-helical transmembrane domain (TMD) and amphiphilic α-helices H2 and H3, connected by flexible linkers. We show that TMD has a preferable orientation in the membrane, while H2 and H3 reside at the membrane surface. Orientation of H2 is strongly influenced by palmitoylation of cysteines Cys40, Cys43 and Cys44. Glycosylation of Asn66 affects the orientation of H3. We also observe that the E protein both generates and senses the membrane curvature, preferably localizing with the C-terminus at the convex regions of the membrane. This may be favorable for assembly of the E protein oligomers, whereas induction of curvature may facilitate budding of the viral particles. The presented results may be helpful for better understanding of the function of coronaviral E protein and viroporins in general, and for overcoming the ongoing SARS-CoV-2 pandemic.

## Introduction

Coronaviruses (CoVs) (order *Nidovirales*, family *Coronaviridae*, subfamily *Coronavirinae*) are enveloped viruses with a positive sense, single-stranded RNA genome of ~30 kb, one of the largest among RNA viruses [1]. CoVs infect birds and mammals, causing a variety of fatal diseases. They can also infect humans and cause diseases ranging from the common cold to acute respiratory distress syndrome. Highly pathogenic human coronaviruses include Severe Acute Respiratory Syndrome (SARS)-CoV, Middle Eastern Respiratory Syndrome (MERS)-CoV and SARS-CoV-2 [2,3]. The outbreaks of SARS-CoV in 2002/3 and MERS-CoV in 2012 led to epidemics. SARS-CoV-2 emerged at the end of December, 2019 causing a pandemic of the coronavirus disease 2019 (COVID-19), which is a novel life-threatening form of atypical pneumonia [4].

Antiviral strategies may be roughly divided into two classes: the measures aimed at prevention of the spread of infections, and treatment of patients who have already contracted the disease. Development of both kinds of strategies benefits greatly from understanding the virus physiology, and in particular the structure and function of viral proteins. Structural biology studies of SARS-CoV-2 have seen rapid progress since the beginning of the pandemic [5]. Whereas most of the key information was obtained using experimental techniques, such as cryoelectron microscopy, X-ray crystallography or NMR, computational approaches were key for some of the findings [6,7]. Among the most notable examples are detailed simulations of dynamics of the most important viral proteins [6,8,9] or even the whole virion [10], early generation of atomic models for all SARS-CoV-2 proteins [11], and high-throughput virtual ligand screening of viral protease inhibitors [12,13].

The genomes of all coronaviruses encode four major structural proteins: the spike (S) protein, nucleocapsid (N) protein, membrane (M) protein, and the envelope (E) protein [14]. The S protein is involved in the host recognition, attachment and cell fusion. The N protein is involved in packaging of the RNA genome and formation of the nucleocapsid. The M protein directs the assembly process of virions through interactions with the other structural proteins and defines the shape of the viral envelope. The E protein is possibly the most mysterious of them since it is associated with the assembly of virions, effective virion transfer along the secretory pathway as well as a reduced stress response by the host cell. Generally, it promotes virus fitness and pathogenesis [15].

Overall, coronaviral E proteins are small, integral membrane proteins of 75–109 amino acids, which have at least one helical transmembrane domain (TMD) and a long amphiphilic region comprising one or two α-helices at the C-terminus [16,17]. SARS-CoV-2 E protein consists of 75 amino acids, and its sequence is 95% and 36% identical to those of SARS-CoV and MERS-CoV E proteins, respectively. Given the sequence identity and the available data, SARS-CoV and SARS-CoV-2 E proteins appear to be very similar in their structure and function, and most of the findings about the former proteins likely apply to the latter as well. The general properties of the SARS-CoV-2 E protein presumably match those of other coronaviral E proteins.

It was shown previously that E proteins may undergo post-translational modifications (PTMs) [16,17], but the role of these modifications is still not fully clear. The prominent examples of other viral proteins that may be also modified by palmitoylation are the coronaviral S protein, haemagglutinin (HA) protein of the influenza virus, Env proteins of retroviruses and filoviruses, and vaccinia virus 37 kDa major envelope antigen (p37) [18–21]. Some results indicate that conserved cysteines SARS-CoV E and, presumably, their palmitoylation are functionally important for stability of the E protein and the overall virus production [22,23]. On the other hand, glycosylation shields viral proteins from recognition by immune system [24,25]. It is closely linked to the protein topology as modification can happen only in the lumen of endoplasmic reticulum. SARS-CoV and SARS-CoV-2 E proteins are predominantly inserted in the membrane with their C-termini in the cytoplasm and are not modified [22,26,27]; a minor fraction can be glycosylated under certain non-native conditions [22,27]. The role of this possible glycosylation of E proteins is not clear [17].

Only a small portion of the E protein expressed during infection is incorporated into the virion envelope; the rest is localized at the intracellular trafficking endoplasmic reticulum (ER)-Golgi region and mostly at the intermediate compartment between ER and Golgi apparatus (ERGIC) [23,26,28–31]. ERGIC is composed of tubulovesicular membrane clusters, with many curved membrane regions [32]. CoVs assemble and bud at the ERGIC, where the E protein may induce membrane curvature or aid in membrane scission [16,17]. Indeed, various recombinant CoVs lacking the gene for E exhibit an aberrant morphology, from which it can be concluded that the function of E is to induce membrane curvature of the viral envelope, thus allowing CoV particles to acquire their characteristic spherical shape and morphology [29,33,34]. E protein was also shown to colocalize and interact with the M and N proteins[35,36] as well as with the N-terminus of nsp3 [37]. It increases the expression of the M and S proteins [38] and, together with M, affects processing, maturation and localization of S [28]. Finally, it can also interact with cellular proteins Bcl-xL [39], PALS1 [40,41] and others such as CWC27, AP3B1, ZC3H18, SLC44A2, BRD2 and BRD4 [42].

In the host membranes, E proteins oligomerize to form ion-conductive pores [43–46] that may be inhibited by hexamethylene amiloride (HMA) and amantadine [47–51]. Similar small proteins (60-120 residues), which oligomerize and form hydrophilic transmembrane pores or channels and disrupt a number of physiological characteristics of the cell, are called viroporins [52]. They are known to contribute to release of infectious enveloped virus particles from infected cells and/or to facilitate the penetration of the viruses into the cell. The most famous representative viroporins of highly pathogenic RNA viruses are human immunodeficiency virus type 1 (HIV-1) Viral protein U (Vpu) protein, hepatitis C virus p7 protein and influenza A virus matrix protein 2 (M2), which are involved in diverse processes such as virus entry, trafficking, assembly, inflammation and apoptosis [52]. The significant contribution of viroporins to the life cycle of viruses makes them a target for therapeutic interventions. In particular, M2 can be targeted by an FDA-approved inhibitor rimantadine [53]. Investigations have shown that SARS-CoV viruses, in which the channel activity is inhibited, were much less infectious and pathogenic [54].

Currently, several experimental structures of the E protein fragments are available [45,48,50,55,56]. In particular, it was shown that the protein contains a TM α-helix and one (when in monomeric form and in detergent, [55]) or two (when in pentameric form, [56]) amphipathic α-helices. Yet, experimental structure of the full-length wild type protein is not available at the moment, and the influence of PTMs on it hasn’t been studied. Moreover, there is little data on behavior of monomeric E protein prior to its assembly into pentameric channels. In the present study, we applied molecular dynamics to study the behavior of monomeric E protein from SARS-CoV-2 and identified the effects of PTMs on the protein behavior. We have also observed that the protein induces curvature in the membranes, and is attracted to the curved regions. These findings may be helpful in development of anti-SARS-CoV-2 medications, and for understanding the function of viroporins in general.

## Results

### Structure of the monomeric E protein

E protein from SARS-CoV-2 is a 75 amino acid-long protein that may be palmitoylated and glycosylated *in vivo*. To assess the overall conformational space available to the protein, we conducted first an extensive coarse-grained (CG) simulation of unmodified E protein, followed by atomistic simulations of unmodified protein and CG simulations of the protein with modifications (Table S1). CG simulations are known to faithfully reproduce the major physicochemical properties of the studied macromolecules while providing a considerable speedup compared to atomistic simulations [57,58].

In accordance with expectations, the simulations revealed that the protein is very flexible with no particular tertiary structure (Figure 1B). Principal component analysis (PCA) shows that the first two components describe most of the structural variation (~64%, Figure 2) and correspond to motions of the helices H2 and H3 relative to each other and TMD near the membrane surface.

**Figure 1.**
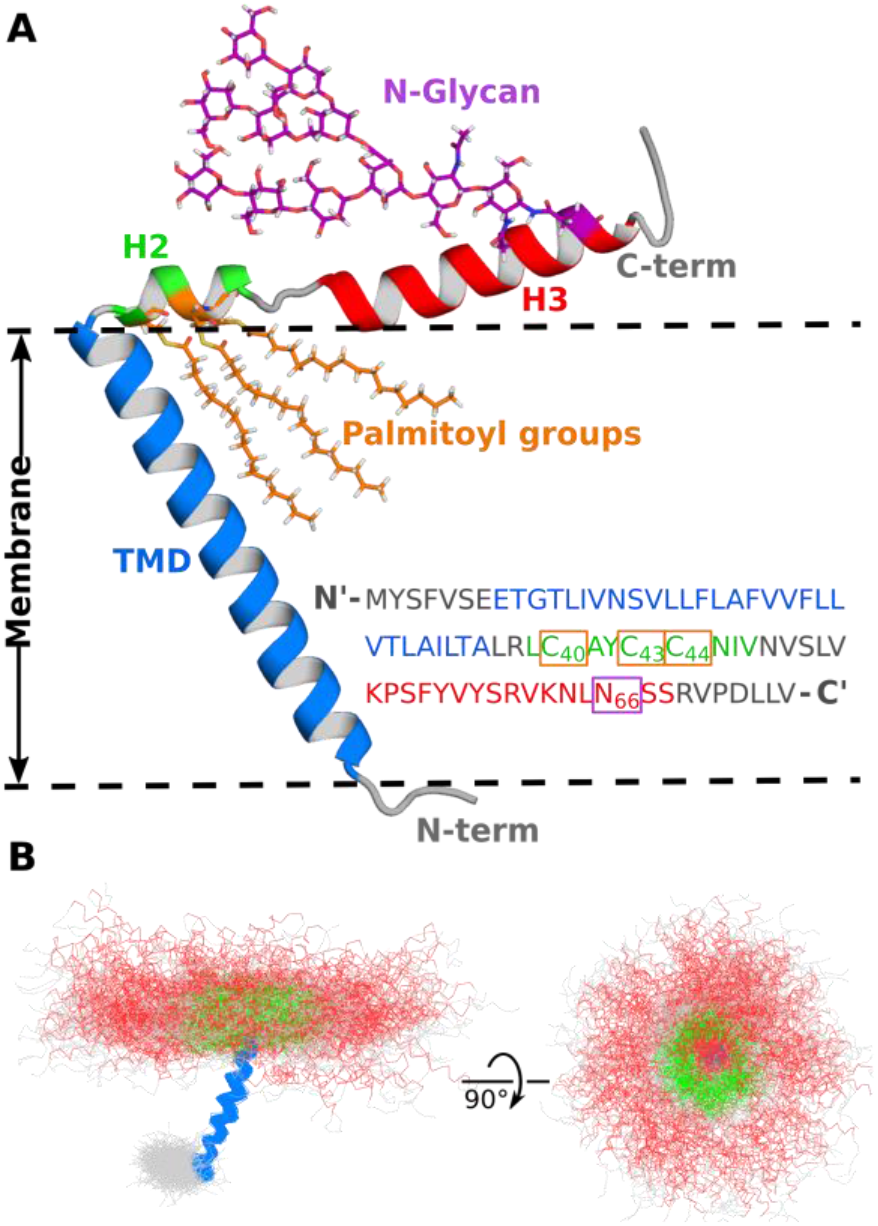
Structure of the SARS-CoV-2 envelope (E) protein monomer with possible post-translational modifications. (A) Schematic model showing a fully palmitoylated and glycosylated E protein. Transmembrane domain (TMD) is shown in blue and amphipathic helices H2 and H3 are shown in green and red. (B) Conformations of the unmodified E protein observed in coarse-grained simulations. Positions of the transmembrane helix were aligned for clarity; helices H2 and H3 are mobile.

**Figure 2.**
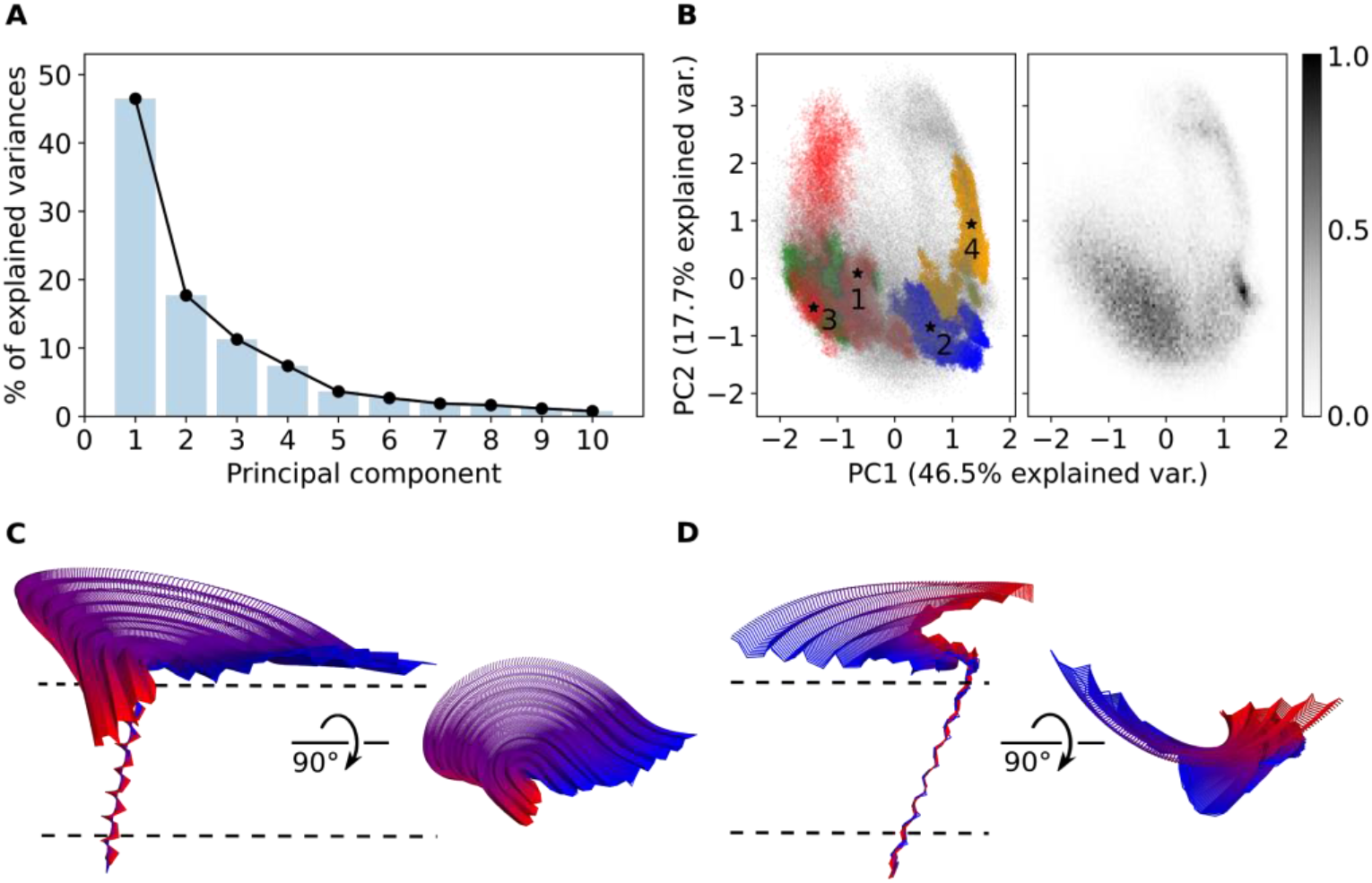
Comparison of the E protein conformations observed in atomistic and coarse-grained simulations using principal component analysis. (A) The scree plot for the top ten PCA eigenvalues. First two components describe ~64% of structural variations. (B) Comparison of the conformational ensembles observed in AA (colored) and CG (gray) simulations projected onto PC1 and PC2. The data for CG simulations only are shown on the right. Starting conformations for AA simulations are shown are labelled with stars. Trajectories from the first, second, third and fourth sets of AA simulations are shown in red, blue, green and orange, respectively. (C) Conformational changes associated with PC1. (D) Conformational changes associated with PC2. The structures are colored from blue to red according to the PC projection value. Approximate membrane position is shown with lines.

Atomistic simulations are considerably more computationally demanding, and thus the exhaustive sampling of the conformational space can take a prohibitively long time. Consequently, we simulated a number of atomistic trajectories starting from representative conformations from the CG simulation. We divided the CG trajectory snapshots into four clusters, and used the centroids of the clusters as the starting structures for atomistic simulations. For each starting structure, we obtained six trajectories of the E protein: three with the protein embedded in the model membrane containing POPC, and three with the membrane mimicking the natural ERGIC membrane (50% POPC, 25% POPE, 10% POPI, 5% POPS, 10% cholesterol). No qualitative differences were observed between the simulations conducted in these membranes. Overall, PCA shows that the atomistic simulations correspond to the CG simulation and display roughly the same conformational space available to the E protein (Figure 2B). Conformations observed in atomistic simulations are shown in the Supporting Figure S1. Atomistic simulations also show that while the secondary structure of the E protein is largely conserved, the amphipathic α-helices H2 and H3 may partially unfold, with H2 being more disordered (Figures 3 and S2). We observed both unfolding and refolding events. Overall, this observation is in agreement with NMR experiments [55,56].

**Figure 3.**
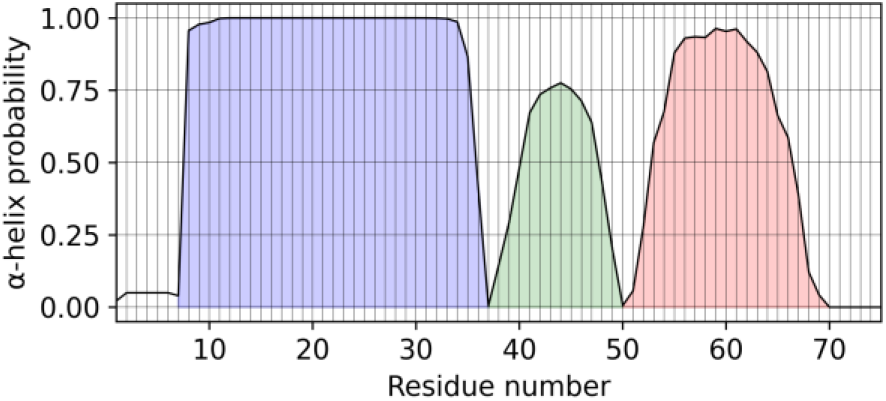
Conservation of the secondary structure of the E protein in AA MD simulations. Average probability of observing the α-helical structure for each residue is shown. TMD remains fully α-helical, whereas H2 and H3 can be sometimes disordered (H2 more often compared to H3).

### Position of the E protein elements relative to the membrane

Figure 4 shows the average positions of the secondary structure elements of the E protein relative to the membrane surface. In all of the simulations, the TMD remained embedded in the membrane. H2 is deeply buried in the lipid headgroup region, whereas H3 is slightly removed from the membrane border, while still remaining in contact with it. In some atomistic trajectories, partial unbinding of H3 from the membrane is observed (Figure S3).

**Figure 4.**
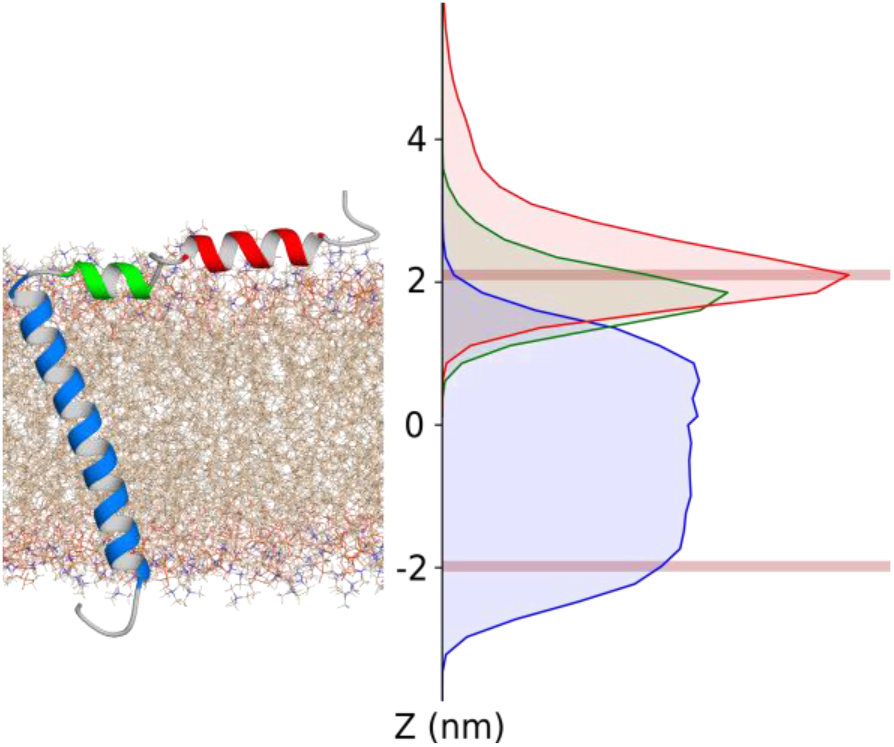
Average positions of TMD, H2 and H3 relative to the membrane in all atom simulations. Average positions of lipid phosphate groups are shown using brown lines. Distributions of TMD, H2 and H3 backbone atoms’ positions are shown in blue, green and red, respectively.

Interestingly, TMD, despite being a single transmembrane α-helix, has a preferable orientation in the membrane (Figure 5). It is tilted at the angle of 25-40° in all of the simulations and has a strong orientational (azimuthal) preference, with phenylalanines Phe20, Phe23 and Phe26 oriented towards the N-terminal side. No robust effects of post-translational modifications on the orientation of TMD were observed (Table 1).

**Table 1.** Average values and standard deviations (σ) of the tilt and rotation angles of TMD and H2 observed in different simulations. See Figures 5 and 6 for definitions.

| Type | System | Lipid | Mean<br>$\alpha, ^\circ$ | $\sigma_\alpha, ^\circ$ | Mean<br>$\beta, ^\circ$ | $\sigma_\beta, ^\circ$ | Mean<br>$\phi, ^\circ$ | $\Delta\phi, ^\circ$ | $\sigma_\phi, ^\circ$ |
| --- | --- | --- | --- | --- | --- | --- | --- | --- | --- |
| CG | No PTM | POPC | 26.9 | 8 | 34.1 | 29.4 | 77.7 |  | 22.5 |
| CG | CYSP40 | POPC | 26.6 | 8.9 | 34.7 | 31.2 | 85.4 | +7.7 | 20.5 |
| CG | CYSP43 | POPC | 26.7 | 8.4 | 32.4 | 29 | 79.3 | +1.6 | 19.7 |
| CG | CYSP44 | POPC | 27.1 | 8.3 | 40.3 | 28.8 | 106.6 | +28.9 | 19.1 |
| CG | CYSP40/43 | POPC | 27.7 | 8.9 | 34.8 | 30.9 | 80.4 | +2.7 | 20.4 |
| CG | CYSP40/44 | POPC | 26.1 | 8.6 | 42.9 | 26.5 | 109.9 | +32.2 | 19.6 |
| CG | CYSP43/44 | POPC | 26.9 | 8 | 36.6 | 31.1 | 96.8 | +19.1 | 17.6 |
| CG | CYSP40/43/44 | POPC | 26.4 | 8.3 | 31.2 | 30.4 | 103 | +25.3 | 17.2 |
| CG | ASNG66 | POPC | 26.2 | 8.5 | 38.7 | 26.9 | 78.6 | +0.9 | 22.5 |
| CG | TMD, no PTM | POPC | 25.5 | 7.7 | 29.4 | 31.4 | - | - | - |
| CG | H2+H3,<br>no PTM | POPC | - | - | - | - | 82.3 | +4.6 | 26.4 |
| AA | No PTM | POPC | 40.2 | 9.9 | 48.1 | 22.6 | 85.3 |  | 49.2 |
| AA | No PTM | Mix | 32.6 | 11.6 | 50.5 | 27.2 | 83.7 |  | 32.6 |

**Figure 5.**
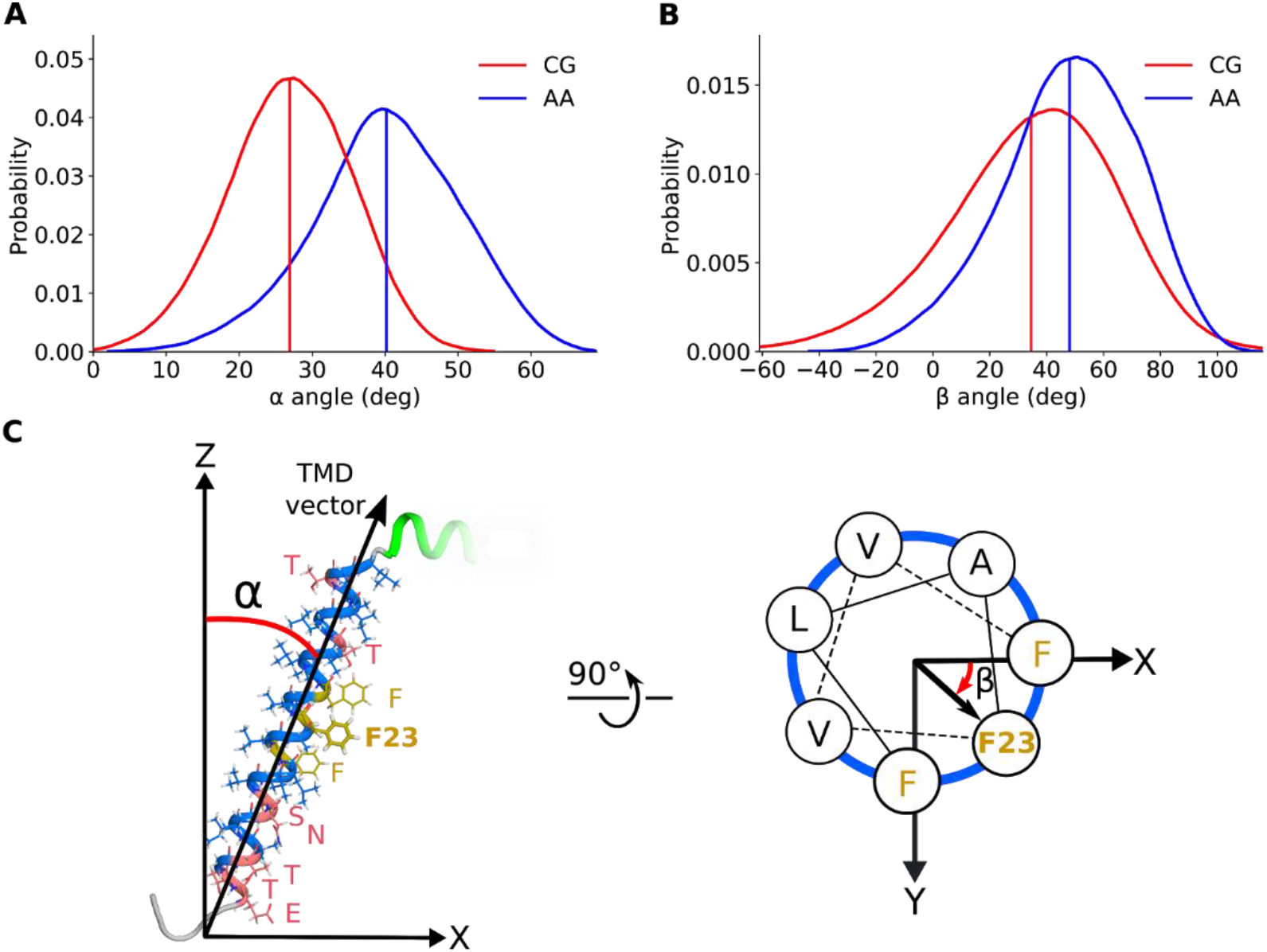
Orientation of TMD in POPC bilayer in coarse-grained (CG) and all atom (AA) simulations. A) Distributions of the tilt angles. B) Distributions of the axial rotation angles. Vertical lines indicate average values. C) Definitions of the tilt (α) and axial rotation (β) angles.

H2, as an amphipathic helix, also has a preferred orientation (Figure 6). Palmitoylation of the three cysteines, Cys40, Cys43 and Cys44, in different patterns changes the physicochemical properties of the helix and leads to its rotation around its axis (Figure 6, Table 1). The strongest effect on H2 orientation is observed when Cys40 and Cys44 are palmitoylated simultaneously: the helix is rotated by ~32.2° relative to its position in the unmodified protein. Other palmitoylated variants have intermediate orientations; the effects of palmitoylation of the three cysteine residues are not additive (Table 1).

**Figure 6.**
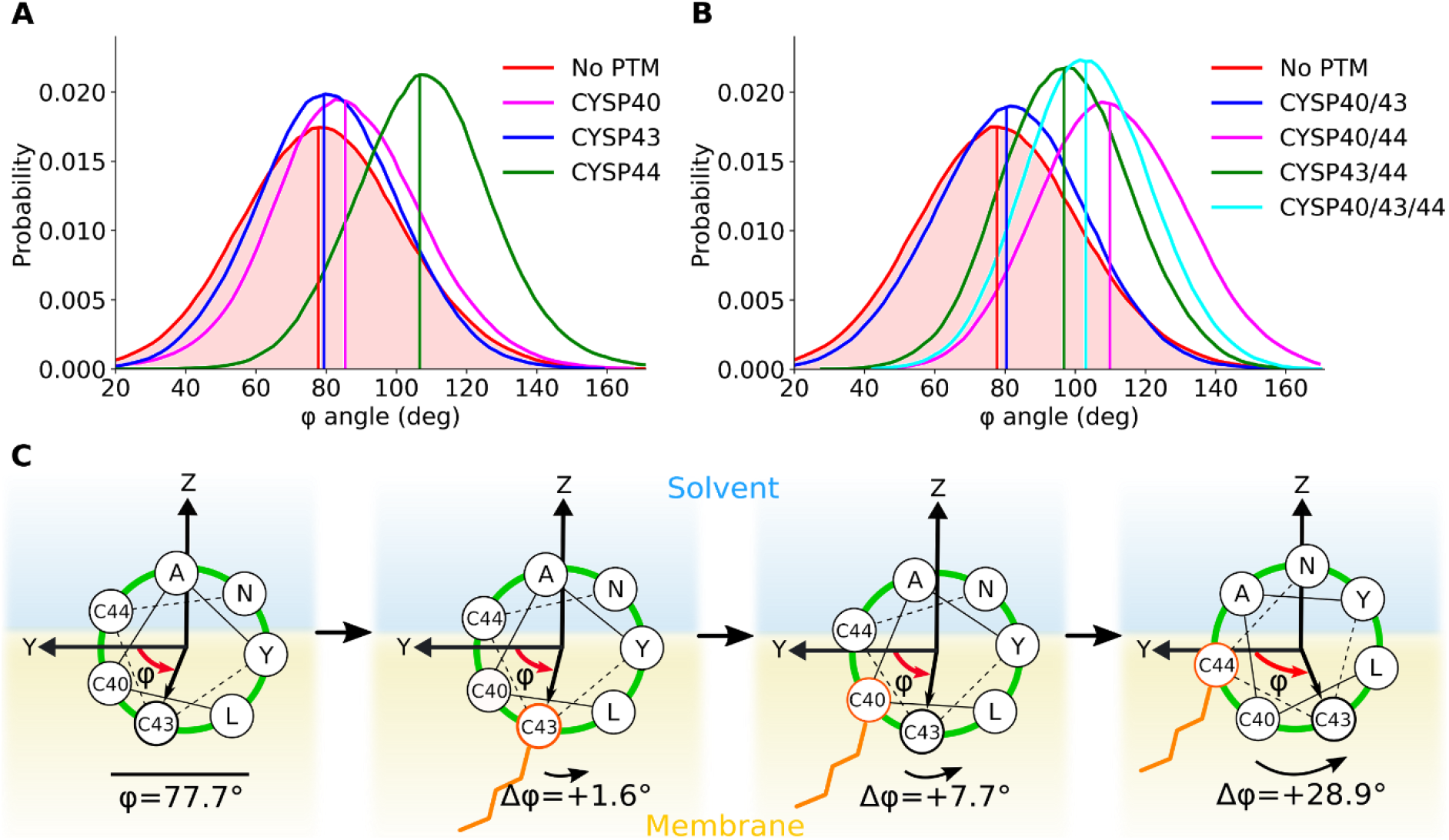
Effects of palmitoylation on orientation of the helix H2 relative to the membrane. Rotation of Cys43 relative to the membrane plane (**Y**) viewed from the N-terminus is analyzed. (A) and (B) Distributions for the Cys43 rotation angles relative to the membrane plane for different PTMs. Vertical lines indicate average values. (C) Schematics showing the H2 orientation with helical wheel projections for selected variants. Palmitoylation affects the orientation of H2 because the respective side chain becomes more hydrophobic.

Position of H3 was not significantly affected by palmitoylation. Yet, glycosylation of Asn66 had a pronounced effect on its dynamics (Figure 7). In the unmodified protein, we observed two major orientations of H3: the first one with the hydrophobic residues Val62 and Leu65 facing the membrane, and the less frequent orientation almost completely opposite to it, with H3 stacking with H2 while being slightly above it (Figure 7). Glycosylation resulted in abolishment of the second orientation, presumably due to the potential steric conflict between the sugar moiety and H2 (Figure 7A,D); the helix was also slightly rotated in the most frequent orientation (Figure 7A,E).

**Figure 7.**
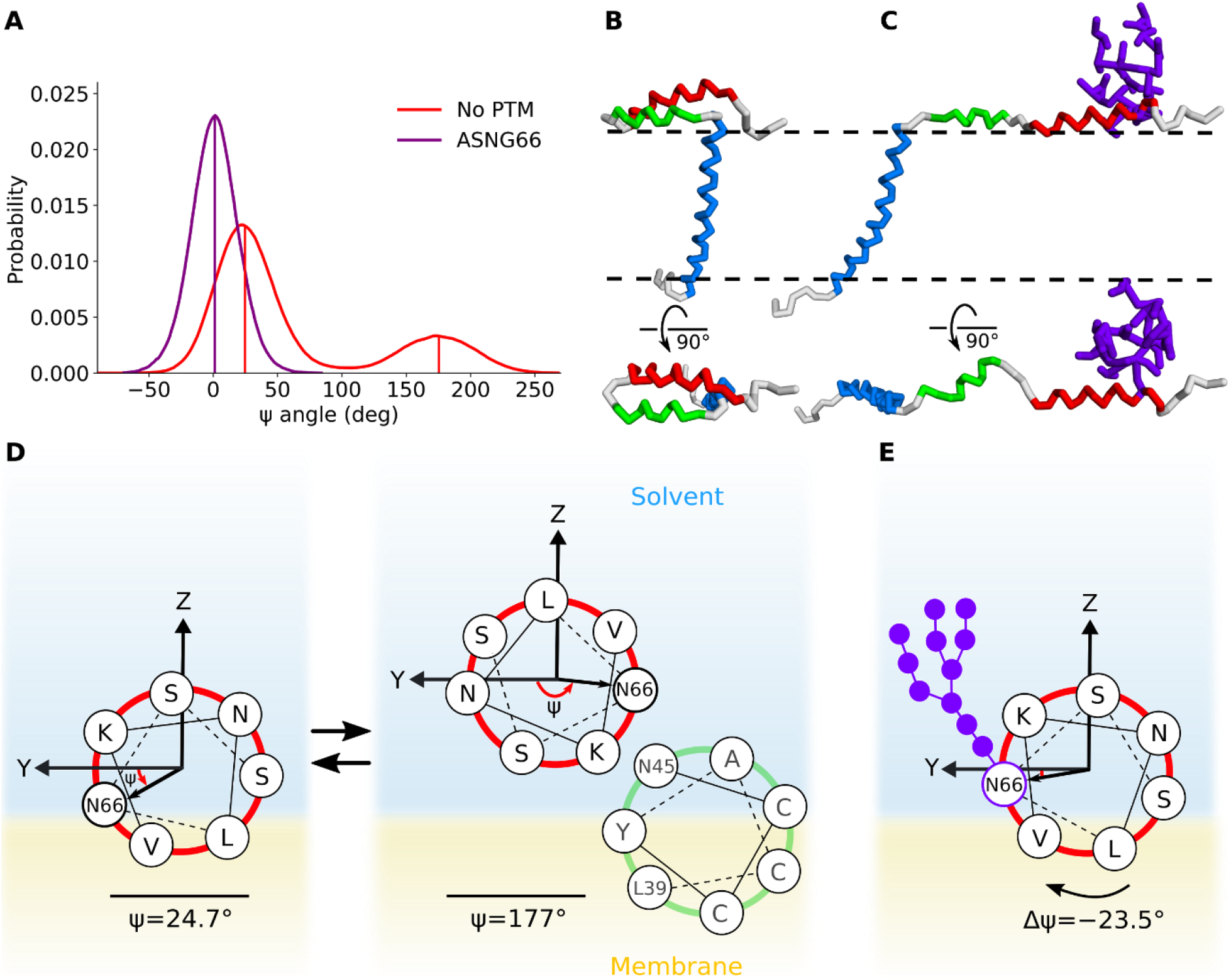
Effect of Asn66 glycosylation on orientation of the helix H3 relative to the membrane in coarse-grained simulations. Rotation of Asn66 relative to the membrane plane (**Y**) viewed from the N-terminus is analyzed. (A) Distributions for the Asn66 rotation angle relative to the membrane plane for unmodified and glycosylated variants. Vertical lines indicate average values. (B) and (C) Representative conformations for ψ ≈ 0° and ψ ≈ 180°. (D) and (E) Schematics showing the H3 orientation with helical wheel projections for unmodified and glycosylated variants. Glycosylation of Asn66 precludes the configuration with ψ = 177°.

### Induction of curvature by the E protein

In all of the conducted simulations, we observed induction of curvature by the E protein: the membrane bends towards the side where the C-terminus is located (Figure 8A). The effect is also observed in larger systems containing four E protein monomers in opposite orientations (Figure S4) and in atomistic simulations (Figure S5). Presumably, the curvature is induced by the amphipathic helices that embed into the adjacent leaflet and expand it.

**Figure 8.**
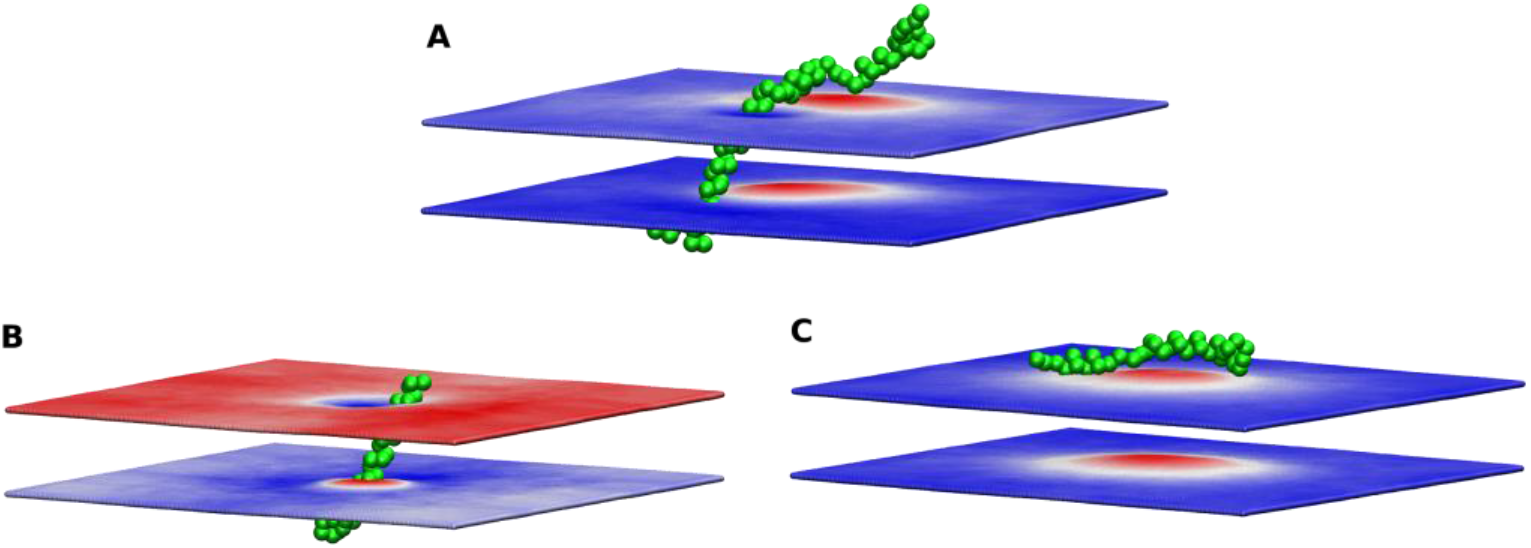
Induction of curvature by the E protein monomer in coarse grained simulations. Upward displacement of each membrane boundary is shown in red, and downward displacement is shown in blue. (A) Induction of curvature by the full-length E protein. (B) Membrane deformation by an isolated TMD. The membrane is thinned around the TMD, but no buckling is observed. (C) Membrane deformation by isolated H2 and H3 helices in coarse grained simulation. The membrane is bent towards the α-helices H2 and H3. Each panel shows an exemplary protein position; positions of the membrane boundaries are averaged over the trajectory length.

To check whether the curvature is indeed induced by H2 and H3, we conducted additional simulations of artificial proteins consisting of only TMD or only H2 and H3 (Figure 8B,C). Isolated TMD was tilted in a way similar to that observed in the simulations of the full-length protein. The membrane was perturbed and thinned near the α-helix (Figure 8B), presumably because of the polar residues on the respective sides (Glu8, Thr9, Thr11, Asn15, Ser16 at the N-terminal side, Thr30, Thr35 at the C-terminal side). Isolated H2 and H3 curved the membrane in the same way as the full-length protein (Figure 8C). Thus, we conclude that the structural elements responsible for curvature induction by the E protein are the amphipathic helices of the C-terminal domain.

### Dynamics of the E protein within curved membranes

Having observed the induction of curvature by the E protein, we were also interested to check whether it has a preferable position in membranes that are already curved, such as the native ER, Golgi and ERGIC membranes, especially during the budding of VLPs. As a test system, we used artificially buckled membranes [59–62]. Irrespective of the starting positions, E protein monomers redistribute in the membranes so that the C-termini localize to the convex regions (Figure 9). The effect was observed both in the membranes buckled in a single direction (non-zero mean curvature, zero Gaussian curvature, Figure 9A) and in the membranes buckled in both directions (positive Gaussian curvature, Figure 9B). Thus, we conclude that the monomeric E protein is curvature-sensitive.

**Figure 9.**
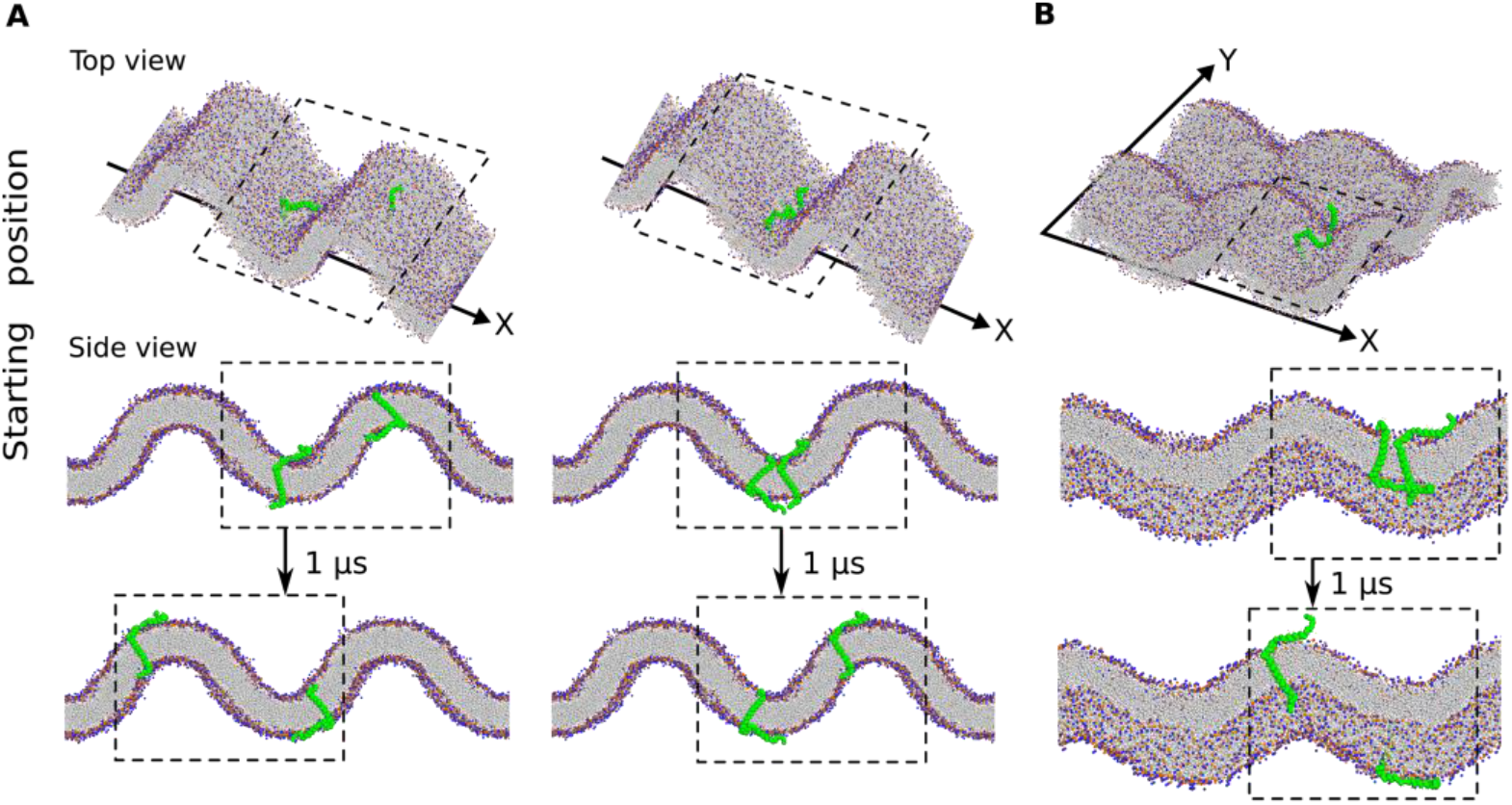
Monomeric E protein partitions into the curved region with the N-terminus localizing to the concave side independent of the starting position. (A) Simulations with the membrane buckled in one dimension. (B) Simulations with the membrane buckled in two dimensions. The dashed box shows the unit cell of the simulation.

## Discussion

Coronaviruses have relatively large genomes harboring tens of different genes. Understanding of their structures could help in development of efficient antiviral measures. Yet some of the CoV proteins are transmembrane, and some contain intrinsically disordered regions [63], at least until they become a part of a larger assembly. E protein has both properties: it has a TM segment, and it lacks tertiary structure in the monomeric form, becoming mostly ordered in the pentameric assembly. Its flexibility and numerous PTMs pose many problems for experimental studies, especially of the monomeric E protein. However, because it is small, and its properties are governed by basic physicochemical principles, it is a good subject for simulations.

Our results show that monomeric SARS-CoV-2 envelope protein has rich conformational dynamics strongly affected by post-translational modifications. The protein is organized as an α-helical TMD and two amphipathic α-helices H2 and H3, flanked by short disordered N- and C-termini. Whereas TMD is rigid and remains α-helical throughout the trajectories, helices H2 and H3 may partially unfold. TMD, H2 and H3 mostly move freely relative to each other, so the monomeric E protein can be considered an intrinsically disordered protein. Yet, all of its α-helices have preferred orientations relative to the membrane.

The TM α-helix of the E protein is relatively long (28 amino acids, ~43 Å), and there is a mismatch between the length of its hydrophobic segment and the thickness of the hydrophobic region of the relevant membranes. Tilting of TM helices is a common mechanism for accommodating such mismatch [64–68]. Accordingly, we find that E protein TMD is tilted at 25-40°, similarly to TMD in pentameric E protein [56] and in other viroporins (Table S2). It also has a preferred azimuthal rotation angle, similarly to WALP peptides [69,70], for which the results obtained with MD simulations were found to correspond well to those obtained in experiments [71]. However, the rotation angle of TMD in a free E protein monomer is opposite to that in the E protein pentamer. This is likely a consequence of the polar residues such as Glu8, Thr9, Thr11, Asn15, Ser16, Thr30 and Thr35 preferably facing the solvent in the monomer and interior of the channel in the pentamer.

One of the most potentially important findings is a strong dependence of the E protein structure on post-translational modifications. Previously, it was shown that palmitoylation is important for E protein stability and overall assembly of VLPs, whereas the role of glycosylation is more elusive [17]. Yet, experimental studies of the effects of PTMs on protein structure are hindered by difficulties in obtaining homogeneous samples with a desired PTM pattern; *in vivo*, palmitoylation is likely to be stochastic [72]. The proteins may be mutated to abolish the particular PTM, however this may also introduce unintended side effects. On the other hand, simulations allow for easier targeted testing of different defined combinations of PTMs.

Previous experimental studies of the E protein dealt with unmodified truncated variants. SARS-CoV E protein construct included residues 8 to 65, a His-tag and a linker; the cysteines were mutated to alanines, Asn66 was missing (Li et al 2014, Surya et al 2018). SARS-CoV-2 construct was even shorter (residues 8-38) and did not include the residues that could be modified (Mandala et al, 2020). Previous computational studies of the E protein also did not focus on the effects of PTM [11,73,74]. In this work, we found that the average orientation of H2 is strongly dependent on palmitoylation pattern, as the acyl chains act as anchors on the respective H2 cysteines and bring them closer to the membrane core. On the other hand, positioning of H3 is affected by glycosylation as the glycan acts as a buoy on H3 and prevents its interaction with H2. However, most or all of the E proteins *in vivo* are probably not glycosylated [27], so the possible role of this modification remains unclear. While we did not study the E protein in pentameric form, we believe that PTMs are likely to elicit effects in the assembled oligomers similar to those that we observe in monomers.

In the last part of our work, we focused on interactions of the E protein with curved membranes. Overall, membrane curvature is an important factor in cell physiology [75], being both generated and sensed by the major membrane constituents: lipids and proteins [76,77]. Some amphipathic α-helices are known to generate curvature by creating the area difference between the two leaflets of the bilayer [78,79] or to act as curvature sensors [80,81].

Along with experiments, molecular dynamics simulations have also been fundamental in studies of curved membranes [57]. Earlier, simulations have been used to study repartitioning of both lipids and proteins in naturally and artificially curved membranes [59–62]; a prominent example is enrichment of cholesterol in the regions with negative curvature [59,61]. Another example is the influenza A М2 protein, for which both experiments and simulations show that its amphiphilic α-helix induces membrane curvature, which is important for VLP budding and membrane scission [82,83], and can act as a curvature sensor [62].

Here, we have found that the SARS-CoV-2 E protein can generate membrane curvature, and this function can be ascribed to the amphiphilic C-terminal domain (α-helices H2 and H2). Such viroporin-generated curvature may stabilize the budding viral particle and promote its formation [75]. We have also found that the monomeric E protein can act as a curvature sensor and localize with the C-terminus at the convex regions of the membrane. Given that the C-terminus of E is oriented towards the cytoplasm [27], the protein is likely to localize at the VLP budding sites and promote VLP budding. Concentration in these curved areas may promote formation of pentameric channels. The assembled channels are also likely to be curvature-sensitive due to their umbrella-like shape [11,56]. On the other hand, E protein is expected to be depleted at the concave inner surface of the VLP, in agreement with experimental data [23,26,28–31].

## Conclusions

Our simulations show that the SARS-CoV-2 E protein in monomeric form has rich structural dynamics. They also highlight the importance of considering the effects of palmitoylation and glycosylation on the protein’s structure. The obtained results are in accordance with experimental observations, while providing a detailed description of the E protein structure. Finally, our work showcases MD simulations as an important complementary technique allowing comprehensive inquiry in the case where the experiment is complicated: when the protein is partially disordered and may be differentially post-translationally modified.

## Acknowledgements

We are grateful to Pavel Buslaev for advices on using the Martini force field. The authors gratefully acknowledge the computing time granted through JARA on the supercomputer JURECA at Forschungszentrum Jülich. I.G. was supported by the Ministry of Science and Higher Education of the Russian Federation (agreement #075-00337-20-03, project FSMG-2020-0003). V.G. and R.A. were supported by the Commissariat à l’Energie Atomique et aux Energies Alternatives (Institut de Biologie Structurale) – Helmholtz-Gemeinschaft Deutscher Forschungszentren (Forschungszentrum Jülich) Special Terms and Conditions 5.1 specific agreement.

## Author contributions

I.G. designed and supervised the project; P.O. and V.G. helped with the project design; A.K. performed simulations with the help of P.O.; A.K. and I.G. analyzed the results and prepared the manuscript; P.O., R.A. and V.G. helped with data analysis and manuscript preparation.

## Additional information

Supplementary information is available.

## Materials and methods

### Model preparation

As a starting structure for simulations of the monomeric SARS-CoV-2 E protein, we used the model prepared by Heo and Feig [11] (https://github.com/feiglab/sars-cov-2-proteins/blob/master/Membrane/E_protein.pdb, accessed on September 1^st^, 2020). Atomistic structure was converted into coarse-grained Martini 3 representation [84] using *martinize*. DSSP [85,86] was used to assign the α-helical secondary structure for TMD, H2 and H3. The CG model was inserted into the POPC lipid bilayer using *insane* [87]. The structure of the monomer with PTMs was constructed manually using PyMOL [88]. For further AA simulations, CG structures were converted to AA using *backward* [89] and inserted into POPC or the native-like mixed bilayer composed of POPC/POPE/POPI/POPS/CHOL in the proportion 10:5:2:1:2 using CHARMM-GUI [90].

In all simulations, the N- and C- termini, residues Lys, Arg, Asp and Glu, and lipids POPS and POPI were charged. The membranes were solvated with water; counter ions were added to neutralize the systems. The simulations were performed using periodic boundary conditions. All systems were energy minimized using the steepest descent method, equilibrated and simulated using GROMACS 2019.5 (AA) and GROMACS 2020.1 (CG) [91]. To generate the buckled membranes, we compressed the bilayers in the *X*-dimension until reaching the desired strain by fixing the box size in these dimensions, while the membrane pressure coupling was turned off in the *X-Y* lateral dimensions.

### Simulation details

CG and AA simulations were conducted using the leapfrog integrator with time steps of 20 and 2 fs, at a reference temperature of 323 and 310 K, respectively, and at a reference pressure of 1 bar. Temperature was coupled using velocity rescale [92] and Nosé-Hoover [93] thermostats with coupling constant of 1 ps^−1^, respectively. Pressure was coupled with semiisotropic Parrinello-Rahman barostat [94] with relaxation time of 12 or 5 ps, respectively.

### CG simulations

CG simulations were conducted using the beta version of Martini 3 force field with nonpolarizable water and optimized parameters for palmitoylated cysteines [95] and glycosylated asparagine [96] where needed. The center of mass of the reference structure was scaled with the scaling matrix of the pressure coupling. The non-bonded pair list was updated every 20 steps with the cutoff of 1.1 nm. Potentials shifted to zero at the cutoff of 1.1 nm and a reaction-field potential with ε_rf_ = ∞ were used for treatment of the van der Waals and electrostatics interactions.

### AA simulations

AA simulations were conducted using the CHARMM36m force field [97]. The covalent bonds to hydrogens were constrained using the LINCS algorithm [98]. The non-bonded pair list was updated every 20 steps with the cutoff of 1.2 nm. Force-based switching function with the switching range of 1.0–1.2 nm and particle mesh Ewald (PME) method with 0.12 nm Fourier grid spacing and 1.2 nm cutoff were used for treatment of the van der Waals and electrostatics interactions. The simulations were performed using JURECA [99].

### Analysis

VMD [100] and in-house scripts were used for analysis of the TMD tilt angle (α) and rotational (azimuthal) angles (β, φ, ψ) of the TMD, H2 and H3. The tilt angle was defined as the angle between the TMD helix axis and the normal to the membrane (axis Z). The rotational angle was defined as the angle between the C_α_ radial vector of a reference residue (Phe23, Cys43, Asn66) and the X-Z plane; the helix was aligned so that its axis was in the X-Z plane (Figures 5-7). We used the Ward’s method from MDTraj [101] to group the dataset into 4 clusters based on pairwise RMSD of coordinates of backbone particles. Density distributions of TMD, H2 and H3 atoms were calculated using the *density* tool from GROMACS. The secondary structure in the E protein was monitored using the *Timeline* plugin (version 2.3) for VMD[100]. PCA was performed on the positions of the C_α_ atoms and backbone particles using the *covar* and *anaeig* tools from GROMACS. Average positions of the membrane boundaries were calculated using *g_lomepro*, version 1.0.2 [102].

