## Supporting information for "Structure and dynamics of the SARS-CoV-2 envelope protein monomer"

**Supplementary Tables S1-S2**

**Supplementary Figures S1-S5**

**Supplementary References**

**Table S1.** List of the simulated systems and their properties.

| № | Type | Protein | Number of lipids (by leaflet) | Number of waters | Number of ions (0.15 mM) | Time, μs | Simulation box size, nm$^3$ |
|---|---|---|---|---|---|---|---|
| 1, 2 | CG | No PTM | POPC (206/221) | 7706 | Na+(84) Cl-(86) | 2, 0.5 | 12×12×11 (rectangular) |
| 3 | CG | CYSP40 | POPC (205/221) | 7711 | Na+(84) Cl-(86) | 0.5 | 12×12×11 (rectangular) |
| 4 | CG | CYSP43 | POPC (206/221) | 7580 | Na+(82) Cl-(84) | 0.5 | 12×12×11 (rectangular) |
| 5 | CG | CYSP44 | POPC (205/221) | 7609 | Na+(83) Cl-(85) | 0.5 | 12×12×11 (rectangular) |
| 6 | CG | CYSP40/43 | POPC (206/221) | 7599 | Na+(83) Cl-(85) | 0.5 | 12×12×11 (rectangular) |
| 7 | CG | CYSP40/44 | POPC (205/220) | 7605 | Na+(83) Cl-(85) | 0.5 | 12×12×11 (rectangular) |
| 8 | CG | CYSP43/44 | POPC (204/221) | 7600 | Na+(83) Cl-(85) | 0.5 | 12×12×11 (rectangular) |
| 9 | CG | CYSP40/43/44 | POPC (205/219) | 7616 | Na+(83) Cl-(85) | 0.5 | 12×12×11 (rectangular) |
| 10 | CG | ASNG66 | POPC (202/221) | 12709 | Na+(139) Cl-(141) | 0.5 | 12×12×15 (rectangular) |
| 11 | CG | TMD, no PTM | POPC (222/219) | 7934 | Na+(88) Cl-(86) | 0.5 | 12×12×11 (rectangular) |
| 12 | CG | H2+H3, no PTM | POPC (207/225) | 7654 | Na+(82) Cl-(86) | 0.5 | 12×12×11 (rectangular) |
| 13 | AA | No PTM (1) | POPC (174/180) | 31286 | Na+(84) Cl-(86) | 0.1×3 | 11×11×12 (hexagonal) |
| 14 | AA | No PTM (2) | POPC (171/179) | 31304 | Na+(85) Cl-(87) | 0.1×3 | 11×11×12 (hexagonal) |
| 15 | AA | No PTM (3) | POPC (169/179) | 30882 | Na+(84) Cl-(86) | 0.1×3 | 11×11×12 (hexagonal) |
| 16 | AA | No PTM (4) | POPC (174/179) | 31104 | Na+(84) Cl-(86) | 0.1×3 | 11×11×12 (hexagonal) |

| | | | | | | | |
|---|---|---|---|---|---|---|---|
| 17 | AA | No PTM (1) | Mixture:<br>POPC (100/100),<br>POPE (50/50),<br>POPI (20/20),<br>POPS (10/10),<br>CHOL (20/20) | 34215 | Na+(151)<br>Cl-(93) | 0.1×3 | 11×11×12<br>(hexagonal) |
| 18 | AA | No PTM (2) | | 32677 | Na+(146)<br>Cl-(88) | 0.1×3 | 11×11×12<br>(hexagonal) |
| 19 | AA | No PTM (3) | | 32243 | Na+(145)<br>Cl-(87) | 0.1×3 | 11×11×12<br>(hexagonal) |
| 20 | AA | No PTM (4) | | 35190 | Na+(155)<br>Cl-(97) | 0.1×3 | 11×11×12<br>(hexagonal) |
| 21,<br>22 | CG | 2×No PTM<br>(X) | POPC (946/950) | 64695 | Na+(710)<br>Cl-(714) | 1 | 25×20×21<br>(rectangular) |
| 23 | CG | 2×No PTM<br>(XY) | POPC (759/761) | 52249 | Na+(573)<br>Cl-(577) | 1 | 21×21×20<br>(rectangular) |
| 24 | CG | 4×No PTM | POPC (3998/3997) | 105411 | Na+(10493)<br>Cl-(10501) | 1 | 50×50×10<br>(rectangular) |

**Table S2.** Properties of the TM helices in experimentally determined structures of single-helical viroporins.

| Protein and virus name | Method | Membrane mimetic | TM residues (length) | Tilt angle | PDB ID | Reference |
|---|---|---|---|---|---|---|
| M2, Influenza A | NMR | DOPC/DOPE | 26-46 (21) | 32° (N-term) 22° (C-term), pH 7.5 | 2L0J | [1] |
| M2, Influenza A | NMR | DMPC | 26-46 (21) | 30° (N-term) 19° (C-term), pH 7.5 | 2KQT | [2] |
| M2, Influenza A | X-ray | OG | 25-46 (22) | ~35° (N-term), pH 7.3 | 3BKD | [3] |
| M2, Influenza A | X-ray | OG | 25-46 (22) | 16° (pH 7.5-8) 31° (pH 6.5) 38° (pH 3-4) | 3LBW | [4] |
| M2, Influenza B | NMR | POPE, POPC/POPG | 6-28 (23) | 14° (pH 7.5) 20° (pH 4.5) | 6PVR, 6PVT | [5] |
| Monomeric Vpu, HIV-1 | NMR | DMPC | 8-25 (18) | 25° | 2N28 | [6] |
| Vpu, HIV-1 | NMR | DOPC/DOPG | 8-25 (18) | 13° | 1PI7, 1PI8 | [7] |
| E, SARS-CoV | NMR | LMPG | 8-35 (28) | ~24° | 5X29 | [8] |

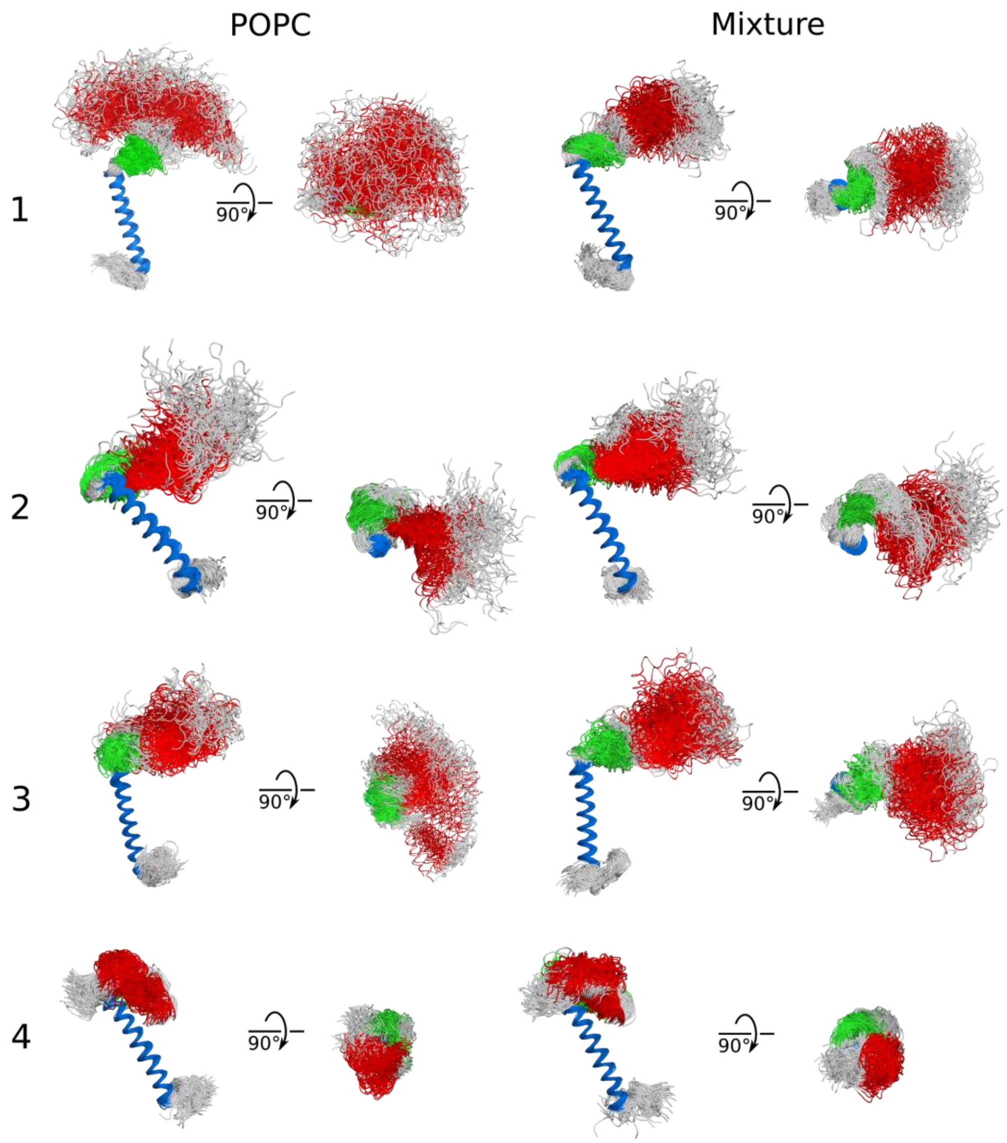


**Figure S1.** Conformations of the E protein observed in all-atom simulations. Positions of the transmembrane helix were aligned for clarity; helices H2 and H3 are mobile.

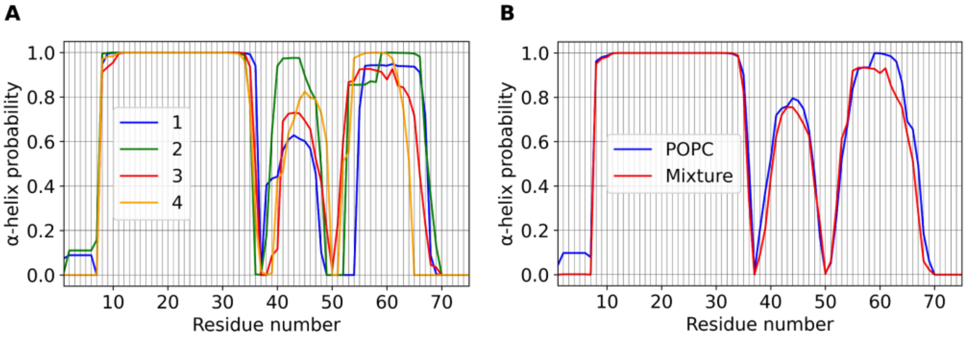


**Figure S2.** Conservation of the secondary structure of the E protein in AA MD simulations. Average probability of observing the α-helical structure for each residue is shown. TMD remains fully α-helical, H2 is sometimes disordered, and H3 is mostly ordered.

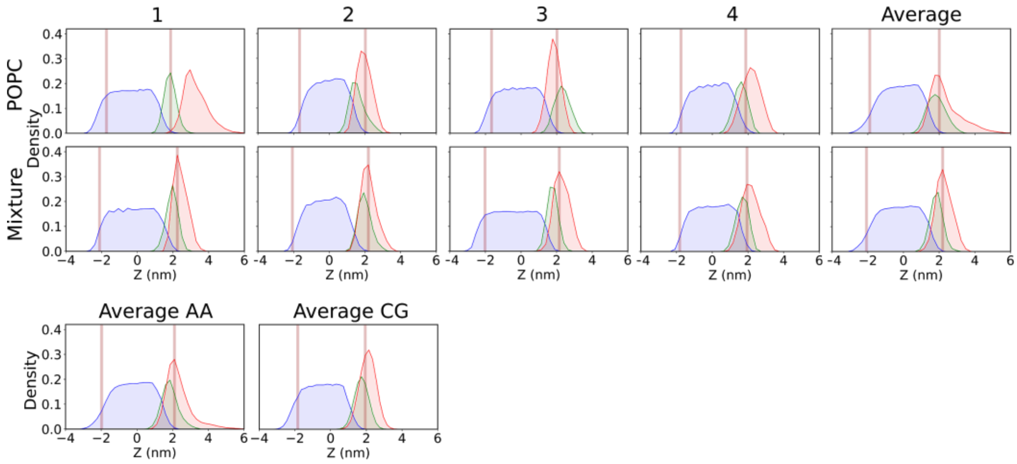


**Figure S3.** Average positions of TMD, H2 and H3 relative to the membrane in CG and AA simulations. Average positions of lipid phosphate groups are shown using brown lines. Distributions of TMD, H2 and H3 backbone atoms' positions are shown in blue, green and red, respectively.

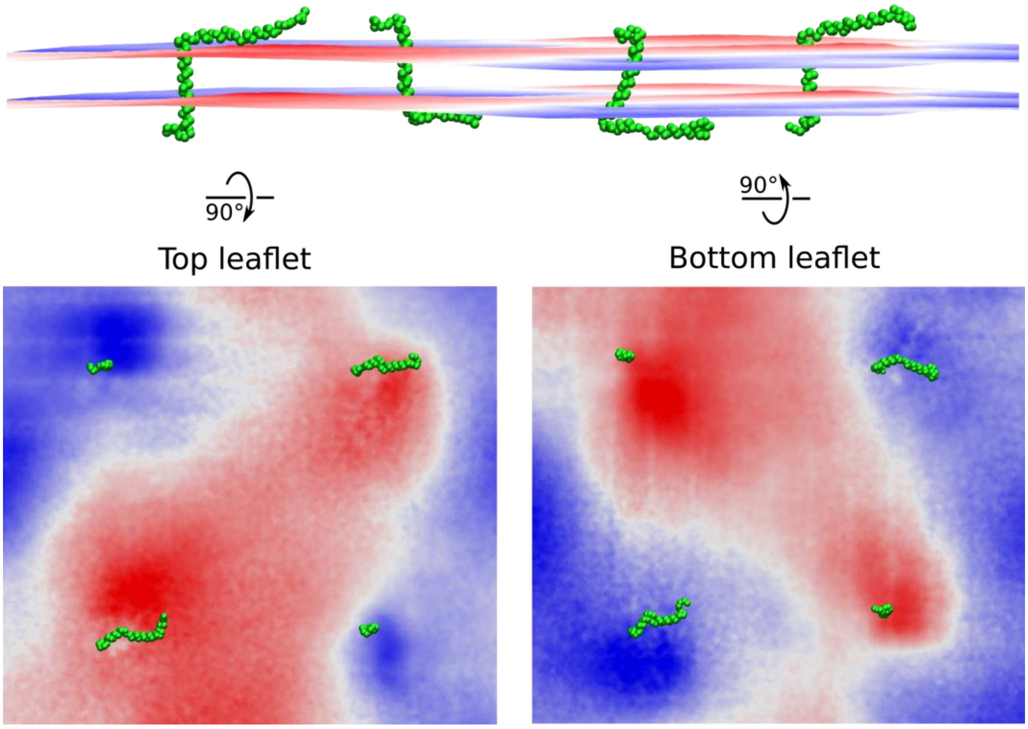


**Figure S4.** Induction of curvature by the E protein monomers in the system containing 4 proteins. (top) Side view of the simulated system. (left) Top monolayer. (right) Bottom monolayer. Upward displacement of each membrane boundary is shown in red, and downward displacement is shown in blue. Each panel shows an exemplary protein position; positions of the membrane boundaries are averaged over the trajectory length of 1 μs.

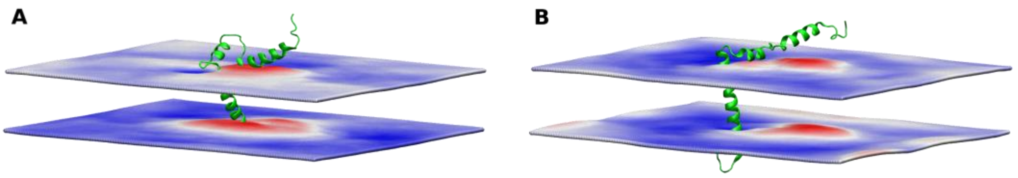


**Figure S5.** Induction of curvature by the E protein monomer in atomistic simulations. Upward displacement of each membrane boundary is shown in red, and downward displacement is shown in blue. (A) Induction of curvature in the POPC membrane. (B) Induction of curvature in the native-like membrane. The membrane is bent towards the α-helices H2 and H3. Each panel shows an exemplary protein position; positions of the membrane boundaries are averaged over the trajectory length.
